# Adaptive diversification and niche packing on rugged fitness landscapes

**DOI:** 10.1101/2022.05.15.492034

**Authors:** Ilan N. Rubin, Yaroslav Ispolatov, Michael Doebeli

## Abstract

Explaining the emergence of diversity and the coexistence of competing types has long been one of the main goals of ecological theory. Rugged fitness landscapes have often been used to explain diversity through the presence of local peaks, or adaptive zones, in the fitness landscape acting as available niches for different species. Alternatively, niche-packing and theories based on limiting similarity describe frequency-dependent selection leading to the organic differentiation of a continuous phenotype space into multiple coexisting types. By combining rugged carrying capacity landscapes with frequency-dependent selection, here we investigate the effects of ruggedness on adaptive diversification and stably maintained diversity. We show that while increased ruggedness often leads to a decreased opportunity for adaptive diversification, it is the shape of the global carrying capacity function, not the local ruggedness, that determines the diversity of the ESS and the total diversity a system can stably maintain.

## 1 Introduction

The adaptive landscape has been a staple of evolutionary thinking since its introduction by Sewall Wright in 1932 (Wright, 1932). Wright proposed that the high-dimensional network of possible gene combinations can be approximately represented by a three-dimensional landscape with two unit-less axes representing genotypes and a third, vertical axis depicting fitness. This theory was later extended to phenotypic trait spaces (e.g., Lande and Arnold, 1983; Whitlock, 1995, 1997; Whitlock et al., 1995), in which the axes represent continuous phenotypic character traits, rather than Wright’s unmeasurable genetic axes, and have largely been evoked interchangeably in subsequent evolutionary theory (e.g., Whitlock, 1995). These landscapes are recognized as a convenient and intuitive tool for visualizing and modelling evolutionary dynamics, despite much debate over the years (Coyne et al., 1997; Wade and Goodnight, 1998). In particular, Wright envisioned rugged fitness landscapes, or landscapes with many local “adaptive peaks” and valleys, and stochastic shifts of a population between peaks as the model for evolutionary dynamics and speciation (Barton and Charlesworth, 1984). For an in-depth discussion of the history of adaptive landscape theory, see Svensson and Calsbeek (2013).

In Wright’s classical fitness landscape, fitness is a static quantity, immutably linked to the genotype or phenotype being modeled and based on characteristics of the given trait or environment. However, it has long been appreciated that the fitness of a given type is affected by the distribution of other types in the population (Clarke, 1979; Maynard Smith, 1982). If fitness is in this way frequency-dependent, the fitness landscape now becomes dynamic and a function of the population present.

While the literature of rugged fitness landscapes explores the dynamics of how a single species adapts to and evolves on that landscape, attempts to explicitly describe adaptive diversification or stable coexistence of multiple types with rugged fitness landscapes remain largely absent. These models are unable to explicitly explain the maintenance of biodiversity if they do not include frequency dependence, a necessary condition for stable coexistence of different types (Chesson, 2000).

In contrast to the rugged fitness landscape model, niche packing and coexistence theory explains the coexistence of many types in a given environment not through the presence of different adaptive peaks, but by the partitioning of a continuous phenotype space into niches (MacArthur, 1970; Roughgarden, 1976). In these models, ecotypes are defined by a continuous phenotype that is used to calculate that type’s carrying capacity (the equilibrium population size of a monomorphic population or the frequency-independent portion of fitness). Phenotype space is then partitioned into distinct niches through frequency-dependent competitive interactions (Leimar et al., 2013a; Szabó and Meszéna, 2006). Classical formulations of the carrying capacity function are unimodal (often Gaussian) or flat. While a limited number of studies consider carrying capacity functions with multiple peaks (Doebeli, 1996; Leimar et al., 2013b; Ranjan and Klausmeier, 2022; Ruxton et al., 2008; Sasaki and Ellner, 1995), the effects of ruggedness in the underlying frequency-independent fitness landscape on frequency-dependent ecological dynamics and multi-species coexistence remain largely unexplored.

Here we use randomly generated rugged fitness landscapes in continuous phenotype space to explore how the emergence and maintenance of biodiversity mediated by negative frequencydependent ecological interactions is affected by the ruggedness of the underlying fitness landscape. Ecology is described by classic Lotka-Volterra dynamics based on competition on a single, continuous phenotype axis and a simple trait substitution process is used to model evolution. Species are defined by a one-dimensional phenotype that controls both the species’ carrying capacity (the frequency-independent part of fitness) and the frequency-dependent competition. The carrying capacity function has “tunable” ruggedness (the number and steepness of local peaks can be controlled), allowing us to explore the effect of increased ruggedness on diversity. In doing so we show that while increased ruggedness does reduce the opportunities for diversification and hinder evolutionary movement in phenotype space, it has little effect on the overall diversity a system can support.

## 2 Model

### 2.1 Tunable rugged landscapes

Here we introduce a model for generating tunable rugged fitness landscapes in continuous trait space. As we consider ecological dynamics including frequency dependence, the landscapes presented here represent the carrying capacity as a function of a continuous phenotype. The carrying capacity can also be thought of as the static portion of fitness, while the frequency dependence due to competition contributes the dynamic portion of fitness that is a function of the species present in the community. The actual fitness landscape is therefore dependent on both the underlying carrying capacity landscape as well as the biological community present and can be measured as the invasion fitness (per capita growth rate of a rare mutant) landscape for any situation. Importantly, carrying capacity is defined as the equilibrium population size of a monomorphic population with a given phenotype and therefore not simply the stated fitness of an arbitrary trait, but a measurable and biologically relevant quantity.

#### 2.1.1 Global carrying capacity function

The carrying capacity landscape can be separated into the underlying global carrying capacity function and the local ruggedness. While the global carrying capacity function can take any form, here we use a quartic function 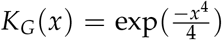 for a given trait *x*. The quartic function has two main advantages. First, the unimodal curve naturally restricts viable phenotypes to an area around the origin (approximately [−2, 2] for the parameterization used here). Second, the relatively flat shape of the curve at the origin (compared to the Gaussian competition function, described below) ensures the presence of a branching point at the origin (when no ruggedness is considered) and therefore the possibility for the coexistence of multiple species (Baptestini et al., 2009; Doebeli, 2011). Additionally, using quartic carrying capacity functions avoids the pitfalls emerging from using a Gaussian carrying capacity function with Gaussian competition (see below), leading to structurally unstable continuous coexistence (Gyllenberg and Meszéna, 2004).

#### 2.1.2 Ruggedness model

To generate the local ruggedness we take a sum of *m* Gaussian distributions along the onedimensional trait axis with equal spacing. The width of each Gaussian is defined by *σ*_*R*_, while the height of each curve, *h*_*k*_, is sampled from a Gaussian distribution: *h*_*k*_ ∼ *𝒩* (*µ* = 0, *σ* = 1). From here on we will refer to *σ*_*R*_ as the period of the local ruggedness. To control the amplitude of the local ruggedness and ensure that there are no negative carrying capacities, the heights are re-scaled so that the deepest “valley” in the local ruggedness has an amplitude of *A*_*R*_. The model becomes:

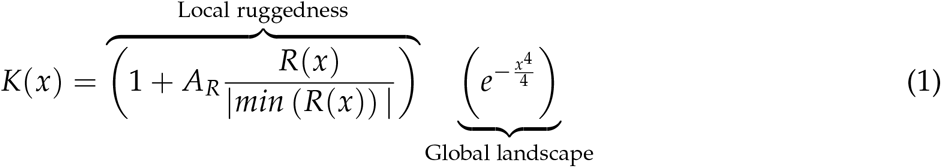

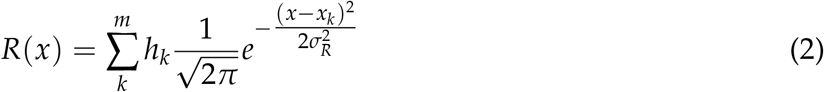

Where

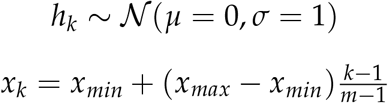

and *x*_*min*_ and *x*_*max*_ represent the extreme range of the modelled phenotype axis.

This process results in a fitness landscape with local ruggedness that is “tunable” by two parameters: *A*_*R*_, the amplitude of the lowest valley relative to the non-rugged carrying capacity function, and *σ*_*R*_, the width of each Gaussian curve that contributes to the ruggedness. Of particular note, the number of Gaussian curves, *m*, used to generate the rugged portion of the landscape only controls the “resolution” of the ruggedness and is uncorrelated to the number of peaks in the final carrying capacity function (Fig. A1). For instance, for the parameters used here (*A*_*R*_ ∈ [0, 1], *σ*_*R*_ ∈ (0, 0.05]), carrying capacity landscapes we generated had between 1 and 76 local peaks, despite each being created as the sum of exactly 250 (*m*) Gaussian distributions spaced equally between -2 and 2 on the one-dimensional trait axis.

A unique carrying capacity landscape is randomly generated for each simulation. For examples of landscapes generated with various values of *A*_*R*_ and *σ*_*R*_ see Figure 1.

**Figure 1:**
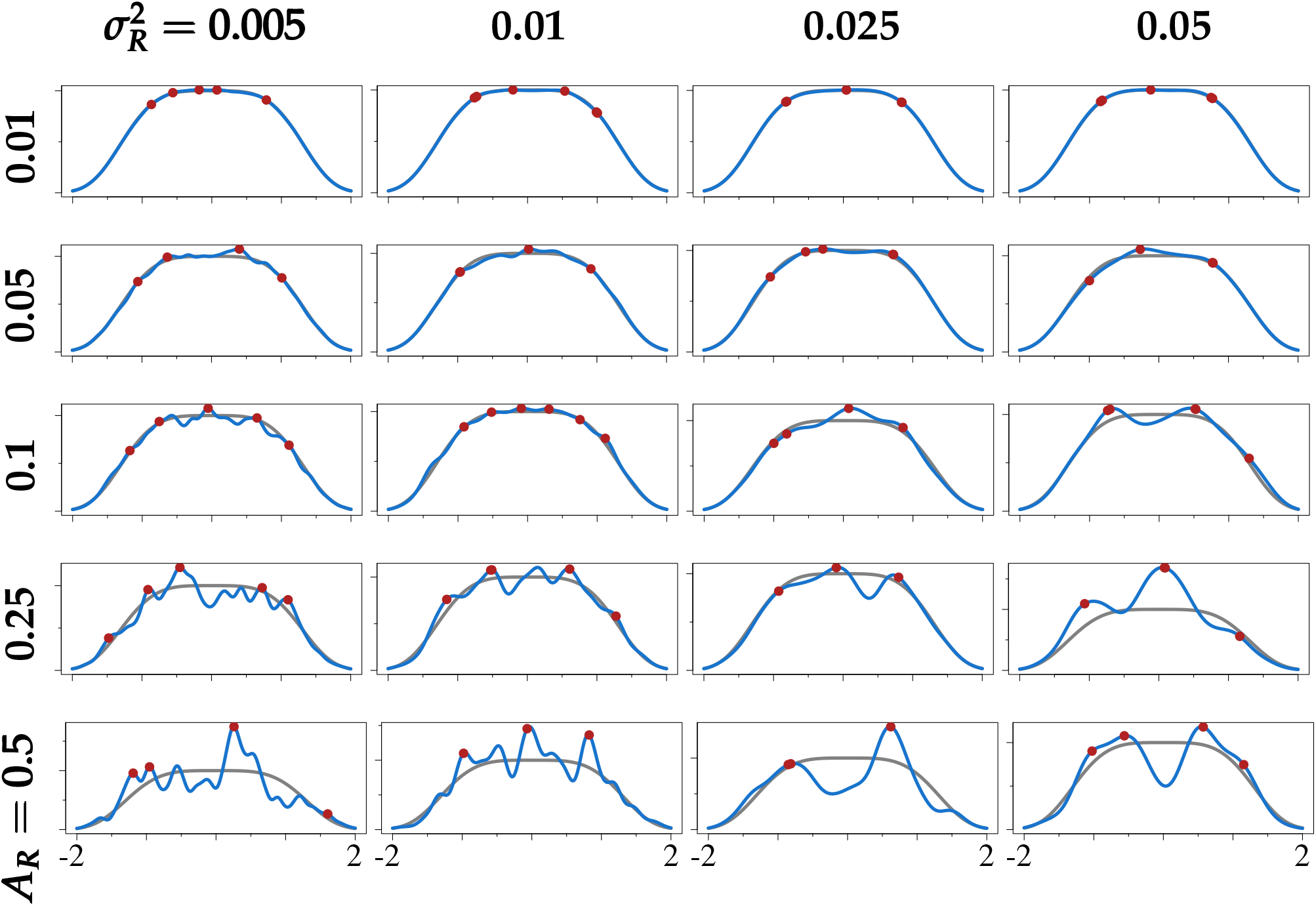
Examples of rugged carrying capacity landscapes with ESS populations. Each rugged carrying capacity landscape is shown in blue, the baseline smooth carrying capacity kernel in grey, and simulated ESS populations in red. Rows denote the amplitude of the local ruggedness (*A*_*R*_ between 1% and 50%) and columns denote square of the period of the local ruggedness (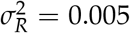 to 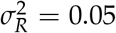). Local ruggedness is generated randomly for each simulation. ESS are simulated by randomly selecting 500 species with phenotype *x* ∈ [−2, 2] and running ecological dynamics until equilibrium (as defined by 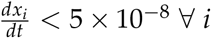 . For pairwise invasion diagrams of the same landscapes please see Fig. A3.

### 2.2 Ecological dynamics

We model ecology using classic Lokta-Volterra dynamics with a single continuous phenotype *x*. All reproduction is assumed to be asexual. The population dynamics for species *i* is defined as:

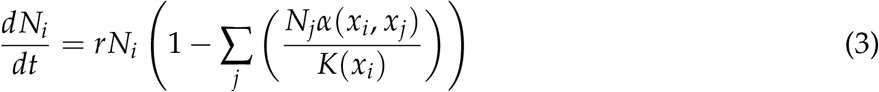

where *r* is the intrinsic growth rate, *N*_*i*_ the population size of species *i, x*_*i*_ its phenotype, *K*(*x*_*i*_) its carrying capacity, and *α*(*x*_*i*_, *x*_*j*_) the competitive effect of species *j* on species *i*. Here we assume competition to be Gaussian, thus the strength of the interaction between two types is symmetric and increases with their phenotypic similarity: 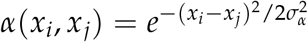. *σ* _*α*_ determines the width of the competition kernel. As *σ*_*α*_ decreases, the interaction strength between two different types decreases.

Ecological dynamics are thus defined by the system of ODEs 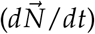 and were numerically integrated using the Runge-Kutta-Fehlberg method. Communities are considered to have reached an ecological equilibrium when *dN*_*i*_/*dt* < 5 × 10^−8^ ∀*i* and any species with a population density *N*_*i*_ < 10^−8^ is considered extinct. All final ecological communities were confirmed to be feasible (i.e., there exists an equilibrium of all species present with strictly positive population sizes) (Grilli et al., 2017) as further confirmation equilibrium was reached. We practically define community diversity as species richness, or simply the number of distinct species present in the community.

As we only consider competitive interactions (as opposed to predator-prey interactions or mutualism, etc.), all communities have a globally stable equilibrium and are therefore indifferent to initial population sizes (Hernández-García et al., 2009; Rubin et al., 2022). Likewise, as we only consider 1-dimensional phenotypes and symmetric competition, evolutionary dynamics always result in a stationary equilibrium (for discussions of non-stationary evolutionary dynamics see Doebeli and Ispolatov, 2014, 2017; Rubin et al., 2021).

### 2.3 Evolutionary dynamics

Evolutionary dynamics are modelled as a trait substitution process. Under the assumption of rare mutations, ecological dynamics are numerically integrated until equilibrium, at which point a new mutant is introduced. Species with population size less than 10^−8^ are considered extinct and removed. Mutants are derived from a randomly chosen parent with probability proportional to their population size and are given a phenotype sampled from a Gaussian distribution centered at the parent phenotype and with variance 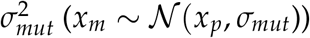. For computational ease, mutant phenotypes were re-chosen if they were closer than *σ*_*mut*_/5 to any resident in the population as populations with very similar species can be exceedingly slow to reach equilibrium. If the mutant has a positive growth rate in the resident population, it is added to the population and the ecological equilibrium is recalculated. Evolutionary simulations were run until 500 consecutive mutants were unable to invade, when an evolutionary equilibrium was declared. All evolutionary simulations were initiated with a single species with a randomly chosen phenotype between −2 and 2. As with the ecological simulations, community diversity is defined as species richness.

The parameterization used here is as follows: 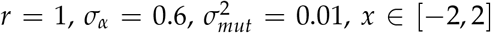, and *m* = 250 (the number of Gaussian curves used to generate local ruggedness). *A*_*R*_ ∼ [0, 1] and 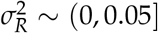.

## 3 Results

### 3.1 Saturated ecological diversity

In order to investigate how ruggedness in the carrying capacity landscape affects the expected number of species that can coexist in a given ecosystem, we conducted numerical simulation experiments saturating a given environment with an over-abundance of species (500 randomly chosen species) and allowing the community to naturally correct down to an equilibrium community. This is a simple, but classic, method of numerically approximating the ESS of a system (e.g., D’Andrea et al., 2019; Rael et al., 2018; Sasaki and Ellner, 1995). The ESS, or evolutionary stable state, of a system is defined as a community that when at ecological equilibrium is completely resistant to invasion (Edwards et al., 2018). This simulation method will inevitably slightly overestimate the ESS (e.g., of the 500 random species, none has the exact phenotype of a species in the ESS – instead, 2 species with similar phenotypes to that ESS species can coexist), but it is still a useful method to compare the diversity different systems can support. After each simulation, the invasion fitness landscape (*x* ∈ [−2, 2]) was calculated. All simulations concluded with less than 4% of the available phenotype space having a positive invasion fitness, verifying this algorithm in approximating the ESS diversity (Fig. A2).

Examples of the landscapes with their final equilibrium populations can be seen in Figure 1. The expected diversity of the ESS of a system with a rugged carrying capacity landscape is relatively unaffected by the period of the ruggedness (*σ*_*R*_) (Fig. 2b). However, for very small amplitude ruggedness (*A* ⪅ 0.02) the resulting diversity is slightly higher than for completely smooth carrying capacity landscapes. For landscapes with larger amplitude ruggedness the diversity of the ESS decreases. In general, systems with low amplitude ruggedness are able to support a more diverse community than those with large amplitude ruggedness, regardless of the period of that local ruggedness.

**Figure 2:**
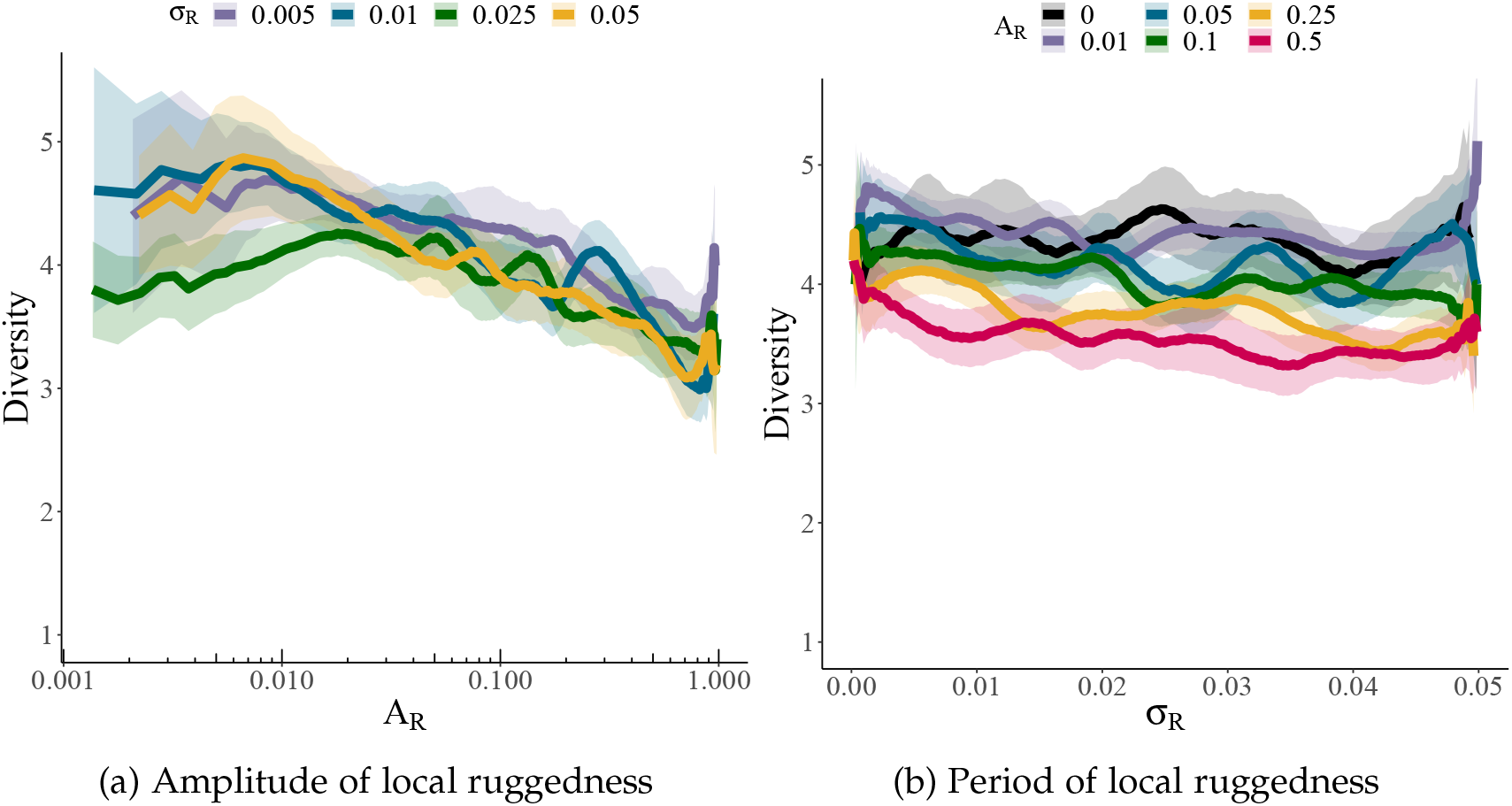
Ecological diversity as a function of the local ruggedness. The final diversity of ecological communities when initiated with a fully saturated initial community (500 species). The lines represent a Gaussian-weighted moving average of the final diversity for a given amplitude (*A*_*R*_) or period (*σ*_*R*_) of randomly generated local ruggedness. The shading represents the 95% confidence interval of the running mean (Gatz and Smith, 1995). The amplitude of the ruggedness is displayed on a log-scale. Between 250 and 500 simulations were run to generate each curve. Values of *A*_*R*_ and 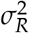 were sampled randomly from [0, 1] and (0, 0.05] respectively. The global carrying capacity is a quartic function and the strength of competition, *σ*_*α*_ = 0.6.

### 3.2 Adaptive diversification

To test the effect of ruggedness on diversification dynamics we placed a single randomly chosen species in systems with the same randomly generated carrying capacity landscapes used previously. Evolution is then allowed to proceed naturally based on the trait substitution process described above. The simulation results in an evolutionarily convergent stable community in which no small-effect mutants can invade. For example communities that result from these evolutionary simulations see Figure A4.

As expected for the competition kernel and carrying capacity used here, in the baseline non-rugged case, evolution always resulted in the 3 species ESS (Fig. 3b) (Rubin et al., 2022). Similarly to the saturated ecological dynamics, small amplitude ruggedness in the carrying capacity landscape resulted in a slightly increased expected equilibrium diversity. For larger amplitudes (*A*_*R*_ ⪆ 0.02), increasing the amplitude of the local ruggedness resulted in decreased diversity at the evolutionary equilibrium. For carrying capacity landscapes with an amplitude near 1, diversification did not occur and the final population remained only the single founding species. While the period of ruggedness (*σ*_*R*_) did not affect the saturated ecological equilibrium diversity, when considering the adaptive diversification from a single species, increasing *σ*_*R*_ leads to increasing diversity. For very small values of *σ*_*R*_ diversification does not occur, regardless of the amplitude of the ruggedness. For landscapes with small amplitude ruggedness, as *σ*_*R*_ increases, the evolutionary equilibrium diversity also increases until it plateaus at a slightly higher expected diversity than the non-rugged baseline case. For landscapes with larger amplitude ruggedness, increasing *σ*_*R*_ also results in increased expected evolutionary diversity, though we did not consider large enough values of *σ*_*R*_ for those curves to reach a maximal level of diversity.

**Figure 3:**
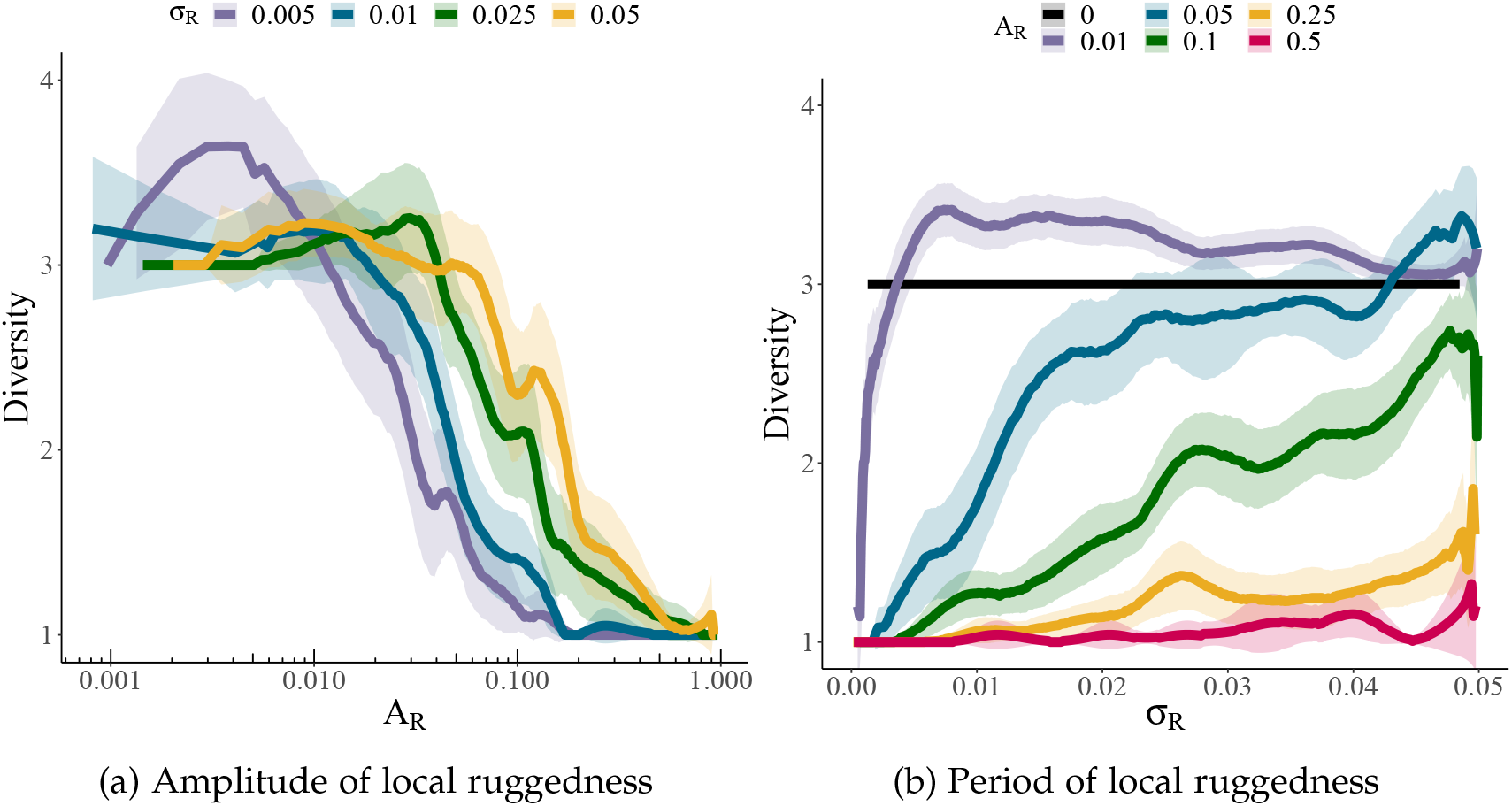
Evolutionary diversity as a function of the width of local ruggedness. The final diversity of ecological communities when initiated with a single, randomly chosen species that is then allowed to evolve. The lines represent a Gaussian-weighted moving average of the final diversity for a given amplitude (*A*_*R*_) or period (*σ*_*R*_) of randomly generated local ruggedness. The shading represents the 95% confidence interval of the running mean (Gatz and Smith, 1995). Of particular note, the line representing *A*_*R*_ = 0 was generated with simulations and is not simply a theoretically calculated value. All simulations with *A*_*R*_ = 0 resulted in 3 species, which is the ESS community for the non-rugged global carrying capacity function. The amplitude of the ruggedness is displayed on a log-scale. Between 250 and 500 simulations were run to generate each curve. Values of *A*_*R*_ and 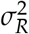 were sampled randomly from [0, 1] and (0, 0.05] respectively. The global carrying capacity is a quartic function with *σ*_*K*_ = 1 and the strength of competition, *σ*_*α*_ = 0.6.

## 4 Discussion

Theories based on rugged fitness landscapes and frequency dependent competition have proposed competing explanations for how diversity is generated and maintained. Rugged fitness landscape theory views local peaks as adaptive zones or available niches waiting to be filled. Evolution by natural selection (and in the asexual populations considered here) is a hill climbing process up the gradient in the fitness landscape to a local peak, at which point adaptation stalls until a stochastic “peak-shift” event. Such an event can be caused by drift in small populations (Wright, 1932), large-effect mutations (Whitlock et al., 1995), or migration (Barton and Charlesworth, 1984) – or more complicated deterministic dynamics like population variance induced peak shifts (Whitlock, 1995) or environmental change causing a shift in the location of the local peaks in phenotype space, viewing rugged landscapes as dynamic seascapes (Mustonen and Lässig, 2009; Simpson, 1953).

Theories of niche packing (Leimar et al., 2013a; MacArthur, 1970; Rubin et al., 2022), evolutionary game theory (Maynard Smith, 1982), and adaptive dynamics (Doebeli, 2011; Geritz et al., 1997) imagine sympatric speciation as an adaptive radiation that is a natural outcome of negative frequency-dependent selection (competition) and selection for individuals to limit their similarity to others in the population. These models are generally built on unimodal carrying capacity functions that represent the frequency-independent, or static, portion of fitness and act as the force of stabilizing selection. Evolutionary dynamics based on frequency-dependent competition initially resemble the same hill-climbing process until the maximum of the carrying capacity landscape is reached. While the maximum is a convergent equilibrium, it may be evolutionarily unstable, i.e., prone to invasion by nearby mutants on either side, resulting in evolutionary branching. The presence of a branching point is determined by the curvatures (as defined by the second derivative) of the competition and carrying capacity functions at that point (or in the one-dimensional case explored here 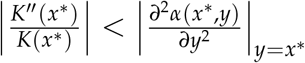 with *x*^∗^ representing the phenotype of evolutionary equilibrium in question) (Dieckmann and Doebeli, 1999; Doebeli, 2011). In this case, there will be disruptive selection at the peak driving the diversification of the population into multiple types. Contrary to scenarios classically imagined through rugged fitness landscape theory, now speciation does not require multiple peaks in a static fitness landscape.

By combining a rugged carrying capacity landscape with frequency-dependent competition we consolidate the intuition and expectations derived from these two bodies of theory. We use a model of classic Lotka-Volterra ecological dynamics with frequency-dependent selection and an underlying rugged landscape for the carrying capacity function. These landscapes have tunable ruggedness, allowing us to control the average curvature and number of local peaks, while leaving the underlying global fitness landscape unchanged.

When considering both frequency-dependent selection and a rugged carrying capacity landscape, the evolutionary dynamics initially are the expected hill-climbing process. However, because selection is frequency-dependent, the actual fitness function does not strictly resemble the rugged carrying capacity landscape. If a local peak and nearby valleys are shallow, the frequency-dependent selection can “flatten” the peak in the dynamically determined fitness landscape, allowing the population to deterministically cross the valley in question and continue to evolve up the global carrying capacity gradient. Alternatively, in systems with very rugged carrying capacity functions (whether because of large *A*_*R*_ or small *σ*_*R*_), this local peak represents a convergent and evolutionarily stable equilibrium and the endpoint of any deterministic evolutionary dynamics. If the landscape has some intermediate level of ruggedness such that the local peak does represent an equilibrium but the carrying capacity is more broadly curved than the competition function at that point in phenotype space, there will exist a convergent stable but evolutionarily unstable equilibrium, i.e., a branching point, and the population will split. This pattern of diversification will continue either until the environment is saturated or adaptation stalls because of a valley in the carrying capacity function being too deep for frequency dependence to flatten, or because of a local peak that is too steeply curved for diversification to occur.

As the landscape becomes more rugged, more local peaks appear in the carrying capacity function and each local peak naturally becomes smaller and with a steeper curvature. Because the stability of any equilibrium in the evolutionary dynamics is dependent on the relative curvatures of the carrying capacity and competition functions at that point, smaller local peaks represent a more stable landscape that retards diversification (Fig. 3). Thus, when the carrying capacity is suitably rugged, the evolutionary dynamics resemble those predicted by Wright’s shifting-balance theory. Directional selection drives the population up the carrying capacity gradient until it becomes trapped on a local peak and must then wait for a stochastic event (whether large mutation, small population-mediated drift, or something else) to cross any nearby area of low fitness (for a discussion of peak-shift dynamics see Coyne et al., 1997; Leimar et al., 2013b; Rouhani and Barton, 1987). With these rugged carrying capacity landscapes, large areas of phenotype space remain invadable but unreachable by mutations of small effect (Fig. A2). When considering frequency dependence these stochastic peak shifts can result in the stable maintenance of multiple types, formalizing the coexistence of many types on rugged fitness landscapes.

Unlike the evolutionary dynamics, for the ecological dynamics of fully saturated communities, the underlying global carrying capacity landscape is the greatest determinant of the level of diversity a system can support, not the nature or severity of the ruggedness of that landscape (Fig. 2). Increasing the amplitude of local ruggedness does result in a less diverse equilibrium community, though the effect is slight compared to the stark effect of ruggedness on diversification dynamics. The total number of local peaks in a landscape has little predictive power for the number of distinct types that the environment can stably support. In very rugged landscapes many of the local peaks in the carrying capacity landscape remain unoccupied and uninvasible, while in less rugged landscapes multiple types may share a single peak.

Interestingly, while increased ruggedness generally depresses diversity in a system relative to the smooth global landscape, small rugged perturbations can actually lead to an increase in community diversity. This can be clearly seen in the small amplitude diversification dynamics (Fig. 3a) and a slight effect in the small amplitude saturated ecological equilibrium (Fig. 2a). This small amplitude ruggedness is not large enough to hinder any evolutionary movement of the population in phenotype space, but does add just enough variation in the carrying capacity to allow diversity to be maintained at phenotypes not included in the smooth carrying capacity ESS.

Of course, a well-known critique of describing evolutionary dynamics on low-dimensional landscapes is that an area of low fitness in one dimension may be circumnavigated through adaptation in higher dimensions (Gavrilets, 1999). While we will not fully comment on higher dimensional landscapes, the general results we focus on here largely stem from the relative curvature of the carrying capacity and competition functions and are thus likely replicated in higher dimensions. However, multi-dimensional evolutionary dynamics are notoriously complex and can result in cyclic and chaotic dynamics (Doebeli and Ispolatov, 2014, 2017; Rubin et al., 2021). How these complex evolutionary dynamics interact with rugged carrying capacity landscapes remains unexplored and could prove insightful to both the peak-shift and adaptive dynamics theories.

### 4.1 Conclusions

Our results affirm that suitably rugged carrying capacity landscapes generate similar diversification dynamics as predicted by peak-shift theory, but not the common narrative that the individual local peaks each represent a specific niche waiting to be filled. Increased carrying capacity ruggedness does impede evolutionary movement in phenotype space and retard diversification. However, the saturated ecosystem diversity is still largely a function of the width of the global niche space compared to the width of the competition, as predicted by niche-packing theory (Doebeli, 2011; Leimar et al., 2013a), rather than the local ruggedness.

## Acknowledgements

YI acknowledges support from FONDECYT (The National Fund for Scientific and Technological Development of Chile) project no. 1200708 (https://www.conicyt.cl/fondecyt/fondecyt-program/).

MD acknowledges support from NSERC (The Natural Sciences and Engineering Research Council of Canada) grant no. 219930 (https://www.nserc-crsng.gc.ca/NSERC-CRSNG/Index_eng.asp)

## 5 Appendix A

### Supplementary figures

**Figure A1:**
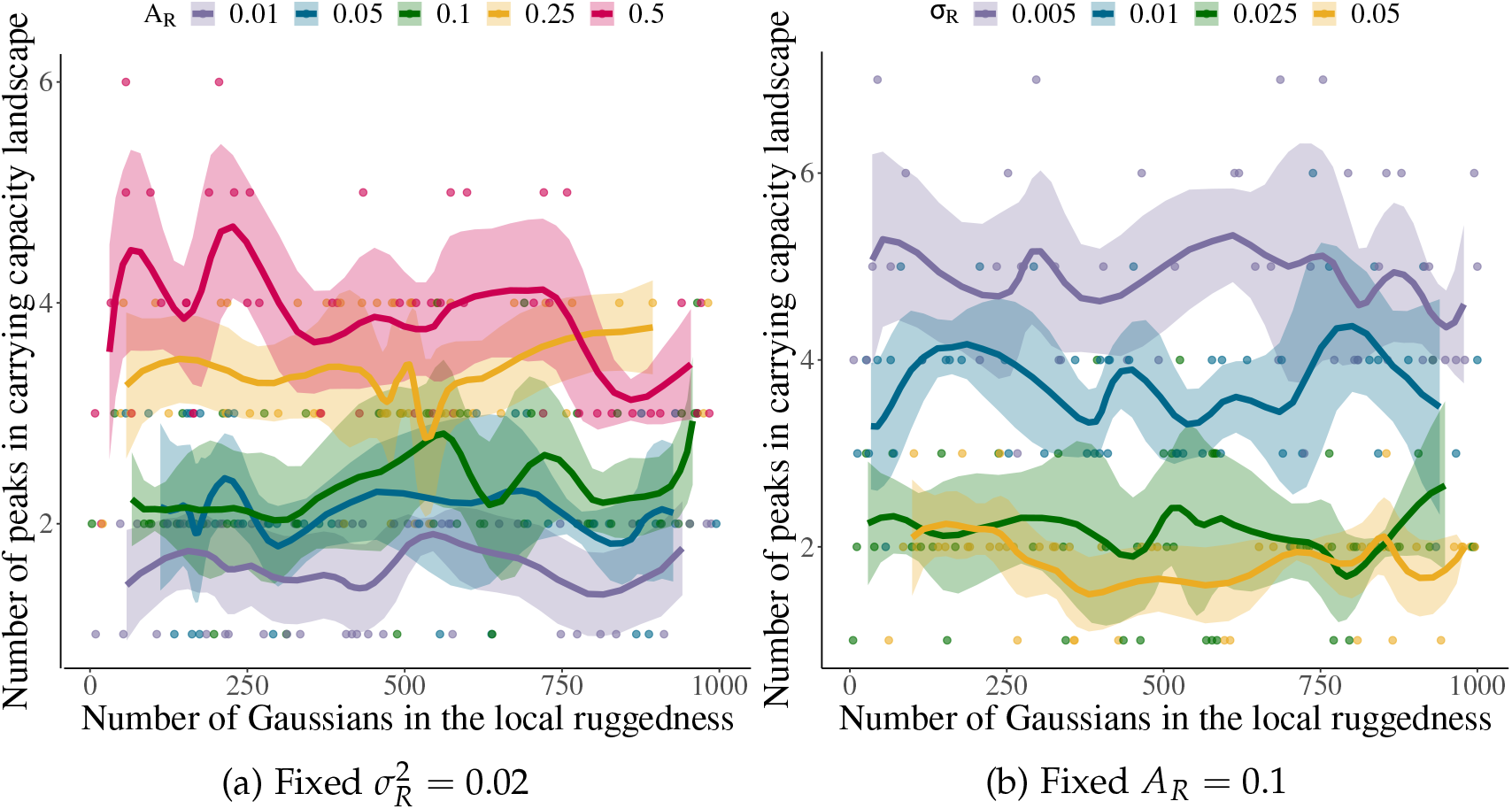
The number of peaks in the carrying capacity landscape. Three parameters used to generate the random ruggedness, the number of Gaussian curves that are laid out at equal intervals between -2 and 2 (*m*), the width of each Gaussian 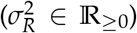, and the amplitude of the lowest valley in the randomly generated ruggedness (*A*_*R*_ ∈ [0, 1]). The number of peaks in the rugged carrying capacity function is relatively indifferent to the number of Gaussian curves used in the generation function (*m*). The number of local peaks was determined numerically by breaking the rugged landscape into 1000 sequential bins. A peak was defined as any bin with a larger carrying capacity than both of its neighbors. For panel A 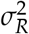 was set to 0.02 and *A*_*R*_ to 0.01, 0.05, 0.1, 0.25, or 0.5. For panel B *A*_R_ was set to 0.1 and 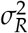 to 0.005, 0.01, 0.025, or 0.05. The number of Gaussians used in the algorithm, *m*, was then chosen randomly between 3 and 1000. Curves represent the running mean with Gaussian weighting and shading the 95% confidence interval of that running mean. 50 landscapes with different values of *m* were used to generate each curve. The global carrying capacity is a quartic function.

**Figure A2:**
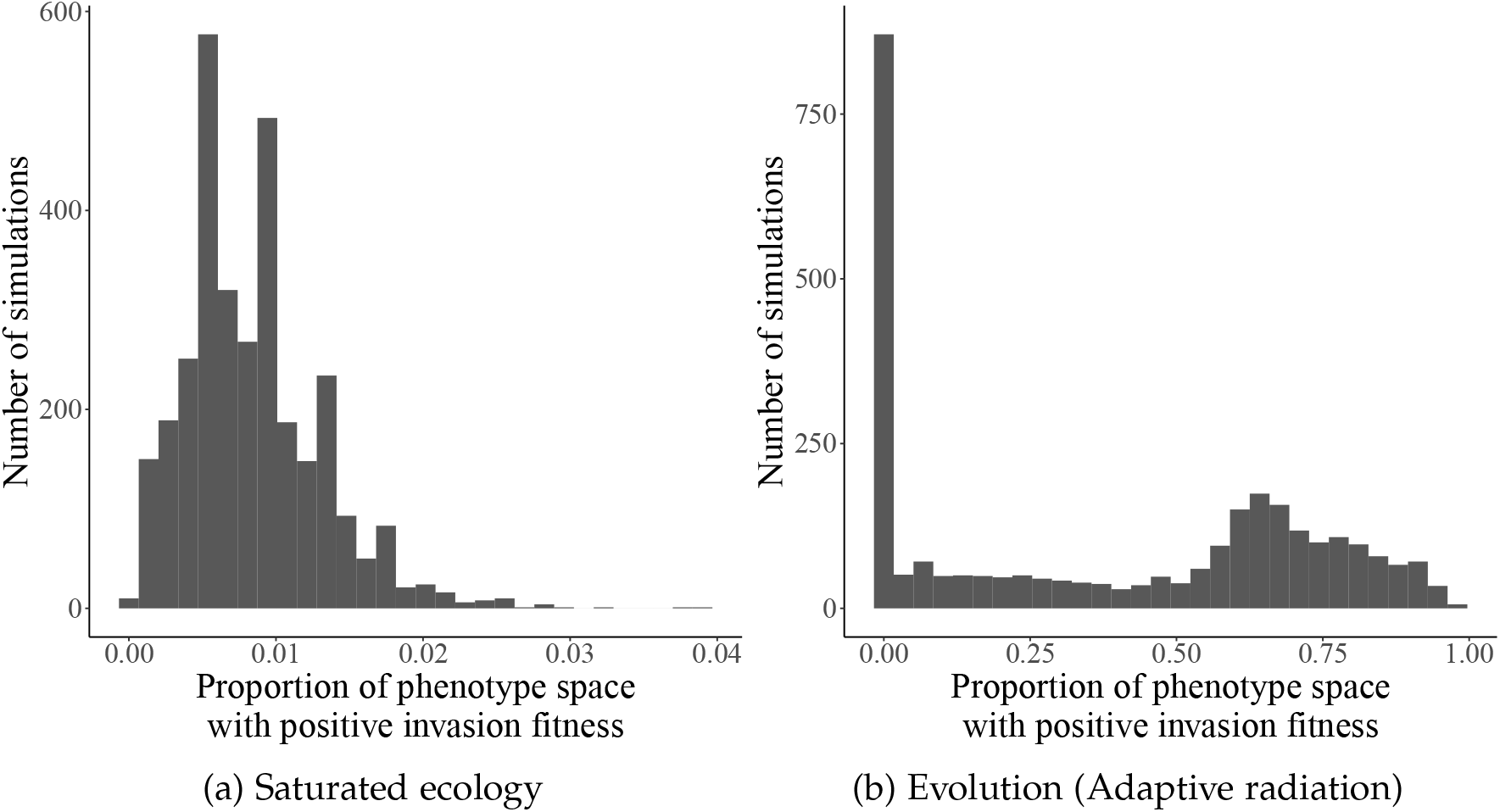
Proportion of phenotype space with positive invasion fitness after ecological saturation. Histogram of the proportion of the phenotype space with positive invasion fitness at the end of the simulation. The ecological simulations starting with a saturated population are shown in panel A and the evolutionary simulations starting with a single species in panel B. Note the x-axis for panel A goes from 0 to 0.06, while the x-axis for panel B is from 0 to 1. Evolutionary simulations with a proportion of invasion fitness equal to 0 represent populations at the ESS. Evolutionary simulations with a very large proportion of the phenotype space open to invasion are due to local ruggedness restricting diversification and evolution in phenotype space.

**Figure A3:**
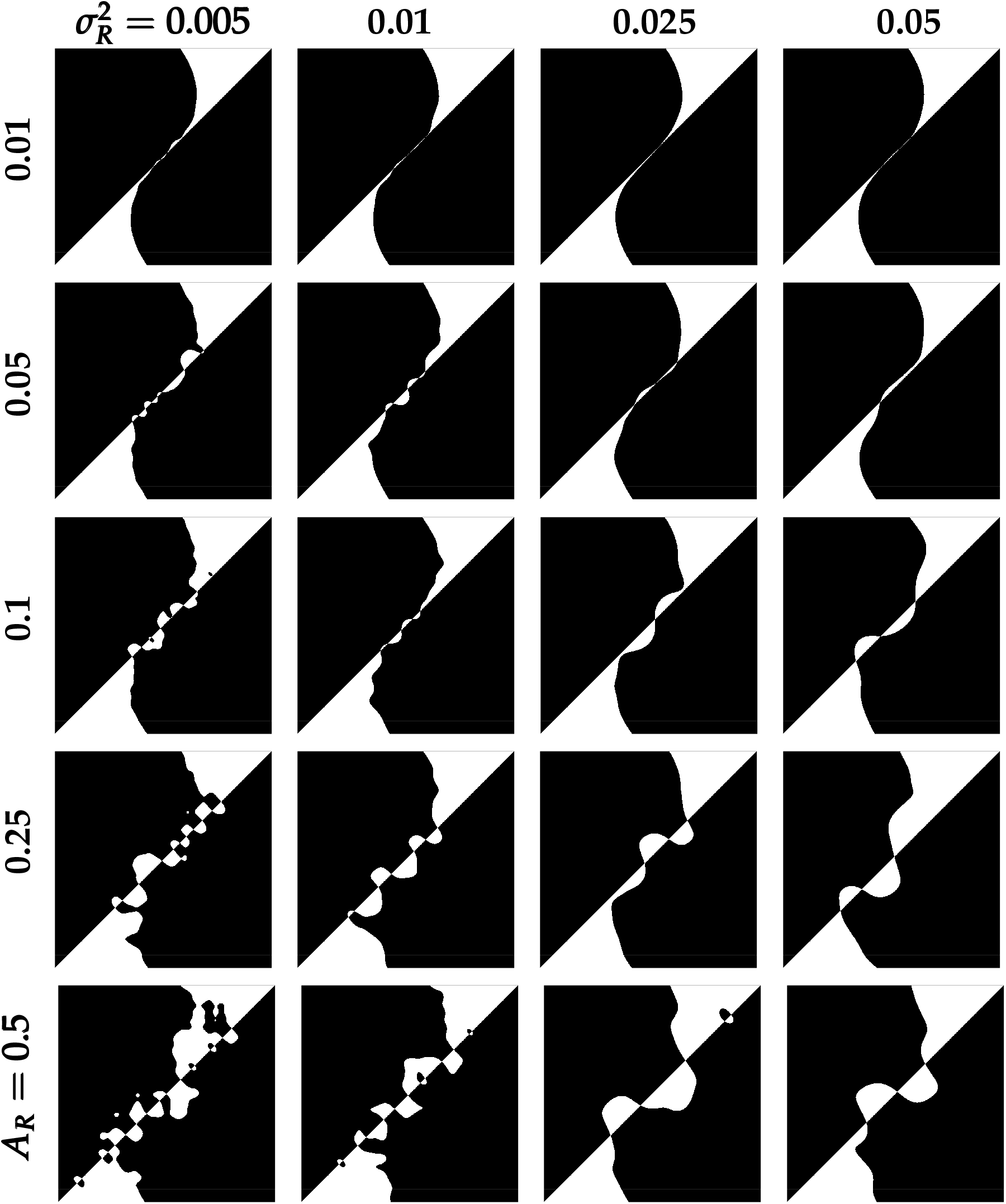
Examples pairwise invasion diagrams for rugged carrying capacity landscapes. Pairwise invasion diagrams for the same landscapes included in figure 1. The x-axis represents the resident phenotype and y-axis the mutant phenotype. The black areas indicate mutants that have a positive growth rate when introduced to a monomorphic population of the resident. If assuming small mutations, the *y* = *x* line is indicative of the evolutionary dynamics. If there is black above and white below *y* = *x*, selective pressure will push the population to evolve to a larger phenotype and vice-versa. Places where the area of positive invasion fitness crosses the diagonal indicate peaks and valleys in the landscape.

**Figure A4:**
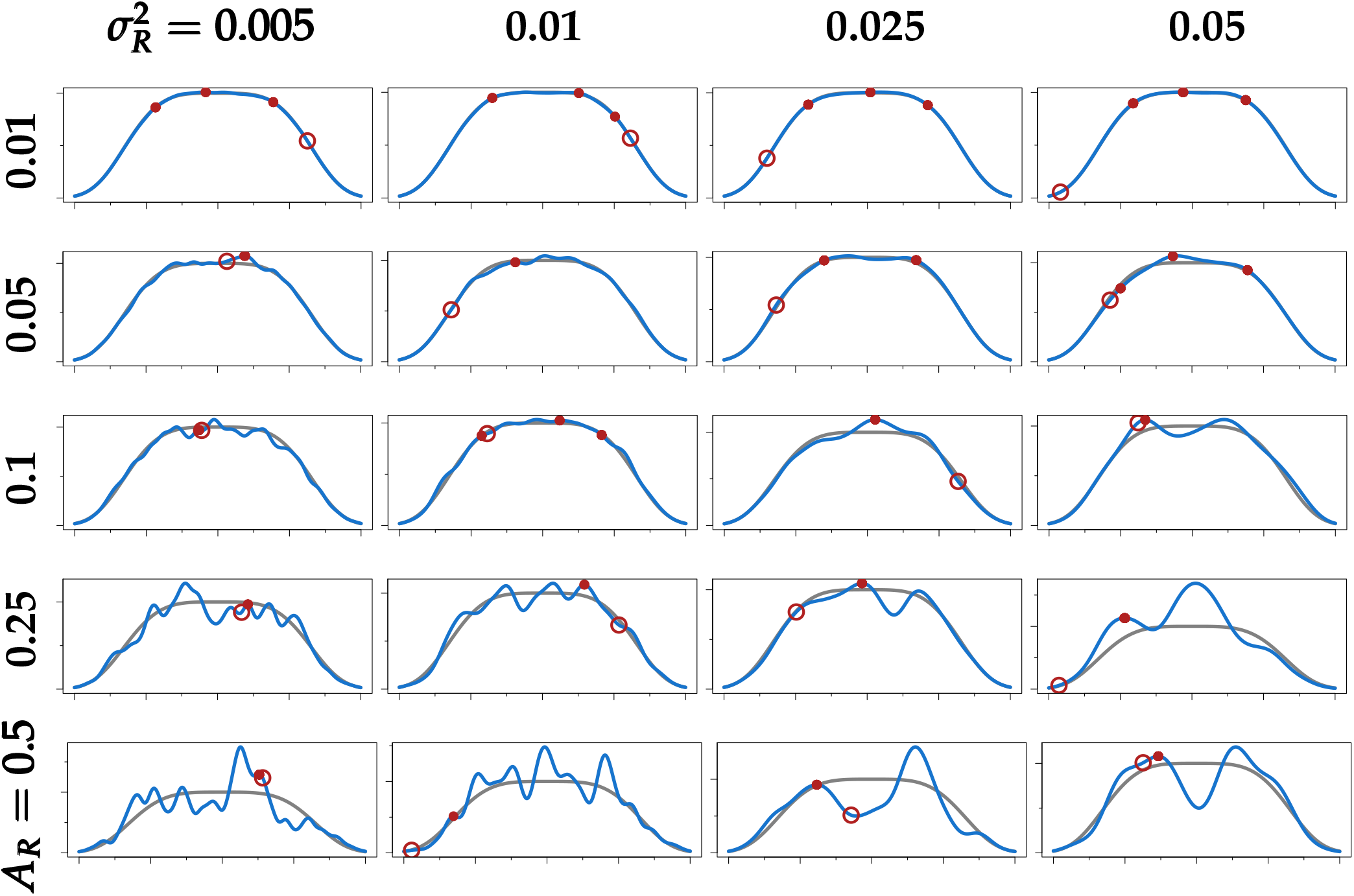
Examples of rugged carrying capacity landscapes with final population after evolution starting from founding species. Each rugged carrying capacity landscape is shown in blue, the baseline smooth carrying capacity kernel in grey, the initial population as a large hollow, red point, and the population after evolution as solid, red points. Rows denote the amplitude of the local ruggedness (*A*_*R*_ between 1% and 50%) and columns denote the period of the local ruggedness (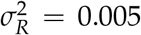 to 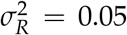). Local ruggedness is generated by adding together 250 Gaussian distributions that are laid out on an equal interval between − 2 and 2, with the width defined by 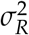 and heights randomly chosen from a Gaussian distribution with *µ* = 1, *σ* = 1. The generated local ruggedness is then re-scaled such that the minimum is equal to the amplitude and then multiplied by the global landscape kernel (*K*(*x*) = *exp*(− *x*^4^/4)) Thus, an amplitude *A*_*R*_ = 0 results in no ruggedness while *A*_*R*_ = 1 means that the lowest valley in the landscape will have a carrying capacity equal to 0. An amplitude *A*_*R*_ = 0.5 means that the lowest valley in the landscape has a carrying capacity equal to half what it would be with a purely smooth carrying capacity function. A single species was randomly selected (*x* ∼ [2, − 2]) as the initial species. Evolution was then simulated through a trait substitution process until 500 consecutive mutants were unable to invade. For pairwise invasion diagrams of the same landscapes please see Fig. A3.

